# Extracellular ATP and CD39 activate cAMP-mediated mitochondrial stress response to promote cytarabine resistance in acute myeloid leukemia

**DOI:** 10.1101/806992

**Authors:** Nesrine Aroua, Margherita Ghisi, Emeline Boet, Marie-Laure Nicolau-Travers, Estelle Saland, Ryan Gwilliam, Fabienne de Toni, Mohsen Hosseini, Pierre-Luc Mouchel, Thomas Farge, Claudie Bosc, Lucille Stuani, Marie Sabatier, Fetta Mazed, Clément Larrue, Latifa Jarrou, Sarah Gandarillas, Massimiliano Bardotti, Charlotte Syrykh, Camille Laurent, Mathilde Gotanègre, Nathalie Bonnefoy, Floriant Bellvert, Jean-Charles Portais, Nathalie Nicot, Francisco Azuale, Tony Kaoma, Jérome Tamburini, François Vergez, Christian Récher, Jean-Emmanuel Sarry

**Author notes:** **Corresponding author:** Jean-Emmanuel Sarry; Inserm, U1037, Centre de Recherches en Cancérologie de Toulouse, F-31024 Toulouse cedex 3, France.

## Abstract

Relapses driven by chemoresistant leukemic cell populations are the main cause of mortality for patients with acute myeloid leukemia (AML). Here, we show that the ectonucleotidase CD39 (ENTPD1) is upregulated in cytarabine (AraC)-resistant leukemic cells from both AML cell lines and patient samples *in vivo* and *in vitro*. CD39 cell surface expression and activity is increased in AML patients upon chemotherapy compared to diagnosis and enrichment in CD39-expressing blasts is a marker of adverse prognosis in the clinics. High CD39 activity promotes AraC resistance by enhancing mitochondrial activity and biogenesis through activation of a cAMP-mediated response. Finally, genetic and pharmacological inhibition of CD39 eATPase activity blocks the mitochondrial reprogramming triggered by AraC treatment and markedly enhances its cytotoxicity in AML cells *in vitro* and *in vivo*. Together, these results reveal CD39 as a new prognostic marker and a promising therapeutic target to improve chemotherapy response in AML.

**SIGNIFICANCE:** Extracellular ATP and CD39-cAMP-OxPHOS axis are key regulators of cytarabine resistance, offering a new promising therapeutic strategy in AML.

## INTRODUCTION

Chemotherapy resistance is the major therapeutic barrier in acute myeloid leukemia (AML), the most common acute leukemia in adults. AML is characterized by clonal expansion of immature myeloblasts and initiates from rare leukemic stem cells (LSCs). Despite a high rate of complete remission after conventional front-line induction chemotherapy (e.g. daunorubicin, DNR, or idarubicin, IDA plus cytarabine, AraC), the long-term prognosis is very poor for AML patients. To date, the 5-year overall survival is still about 30 to 40% in patients younger than 60 years old and less than 20% in patients over 60 years. This results from the high frequency of distant relapses (50 and 85% for patients younger and older of 60 years of age, respectively) caused by tumor regrowth initiated by chemoresistant leukemic clones (RLCs) and characterized by a refractory phase during which no other treatment has shown any efficacy thus far (1, 2). Even with recent efficient targeted therapies that are FDA-approved or under clinical development, therapy resistance remains the major therapeutic barrier in AML. Therefore, understanding the molecular and cellular mechanisms driving chemoresistance is crucial for the development of new treatments eradicating RLCs and to improve the clinical outcome of these patients.

The biological basis of therapeutic resistance (drug efflux, detoxification enzymes, inaccessibility of the drug to the leukemic niche) currently represents an active area of research. However, the molecular mechanisms underlying AML chemoresistance are still poorly understood, especially *in vivo*. It is nevertheless increasingly recognized that the causes of chemoresistance and relapse reside within a small cell subpopulation within the bulk of leukemic cells. Supporting this idea, clinical studies have shown that the presence of high levels of CD34^+^CD38^low/-^CD123^+^cells at diagnosis correlates with adverse outcome in AML patients in terms of response to therapy and overall survival (3, 4). Consistent with these data, Ishikawa and colleagues (5) have observed that this population is also the most resistant to AraC treatment *in vivo*. As a first step towards successful therapeutic eradication of these RLCs, it is now necessary to comprehensively profile their intrinsic and acquired characteristics. We have recently established a powerful preclinical model to screen *in vivo* responses to conventional genotoxics and to mimic the chemoresistance and minimal residual disease observed in AML patients after chemotherapy (6). Accordingly, we have fully analyzed the response to chemotherapy of leukemic cells in AraC-treated AML patient-derived xenograft (PDX) mouse models. Surprisingly, we have found that AraC treatment equally kills both cycling and quiescent cells and does not necessarily lead to LSC enrichment *in vivo*. However, we observed that AraC chemoresistant leukemic cells present elevated oxidative phosphorylation (OxPHOS) activity and that targeting mitochondrial oxidative metabolism with OxPHOS inhibitors sensitizes resistant AML cells to AraC (6, 7). Consistent with our findings, several groups have also demonstrated that essential mitochondrial functions contribute to resistance to multiple treatments in other cancer types (8–11).

Hyperleukocytosis is a clinical condition observed in AML patients, which may lead to life-treatening complications such as leukostasis and is associated with a higher risk of relapse. Importantly, this condition is sustained by several mediators of inflammation, which were also reported to contribute to chemoresistance in AML (12–14). Supporting this idea, a recent study reported that inhibition of the inflammatory chemoresistance pathway with dexamethasone improved AML patient outcome (15). In line with these observations, recent work from our group (6) has highlighted a gene signature involved in the immune and inflammatory response after AraC treatment of PDX models *in vivo*. Amongst immune response mechanisms, the adenosine signaling pathway is one of the most prominent in cancer. CD39/ENTPD1 (ectonucleoside triphosphate diphosphohydrolase-1) is a member of the family of ectonuclotidases present on the outer surface of cells and a key component of the adenosine signaling pathway. Together with CD73, CD39 catalyzes phosphohydrolysis of extracellular adenosine triphosphate (eATP) and adenosine diphosphate (ADP) to produce adenosine, a recognized immunosuppressive molecule (16, 17). Therefore, CD39 has a critical role in tumor immunosurveillance and inflammatory response. Furthermore, although other nucleoside triphosphate diphosphohydrolases (NTPDases) exist, CD39 appears to be the main NTPDase in T lymphocytes and regulatory T cells (18). Recent lines of evidence have revealed high expression and activity of CD39 in several blood and solid tumors (such as head and neck cancer, thyroid cancer, colon cancer, pancreatic cancer, kidney cancer, testis cancer, and ovarian cancer), implicating this enzyme in promoting tumor growth and infiltration (19) and CD39 blockade was recently shown to enhance anticancer combination therapies in preclinical mouse models of solid tumors (20). Furthermore, CD39 is frequently detected in primary tumor cells, including AML blasts, cancer-derived exosomes and tumor-associated endothelial cells. Notably, CD39 was reported to contribute to the immunosuppressive microenvironment in AML (21), while extracellular nucleotides (ATP, UTP) can inhibit AML homing and engraftment in NSG mice (22).

In the present study, we employed computational analysis of transcriptomic datasets obtained from PDX models treated with AraC and from primary patient samples to identify new druggable and relevant cell surface proteins specifically expressed by RLCs. Among these genes, we uncovered CD39/ENTPD1 and confirmed that CD39 expression and activity are increased in residual AML cells post-chemotherapy *in vitro*, *in vivo* and in the clinical setting. Herein, we have also shown that high CD39-expressing resistant AML cells rely on an enhanced mitochondrial metabolism and are strongly dependent on the cAMP-PKA-PGC1α axis. Accordingly, targeting CD39 markedly enhanced AraC cytotoxicity in AML cell lines and primary patient samples *in vitro* and *in vivo* through the inhibition of mitochondrial OxPHOS function and this effect could be mimicked by inhibition of the PKA pathway. Overall, this work shows that the mechanism of resistance to AraC involves CD39-dependent crosstalk between the energetic niche and AML mitochondrial functions through the CD39-cAMP-PKA signaling axis.

## RESULTS

### Enhanced CD39/ENTPD1 expression and activity are involved in early resistance to cytarabine in AML

In order to identify new potential therapeutic targets involved in the onset of AraC resistance *in vivo*, we analyzed a previously identified signature of 68 genes that are significantly upregulated in residual AML cells from PDXs upon AraC treatment *in vivo* ((6); GSE97393). Bioinformatic analysis of this specific gene signature showed an enrichment in several key cancer and immune response signaling pathways, including 8 genes involved in the inflammatory response (Supplementary Fig. S1A). As inflammation has been previously shown to play a critical role in the development of chemoresistance and to be linked to a poor prognosis in AML (15), we focused on this latter group of genes. Within this subset, we identified five genes encoding plasma membrane proteins sensitive to existing inhibitors, thus representing relevant druggable targets of RLCs *in vivo.* Importantly, two of these genes, ecto-nucleoside triphosphate diphosphohydrolase-1 ENTPD1 (CD39) and fatty acid translocase CD36, were specifically overexpressed in AML cells compared to normal HSCs, highlighting their potential as therapeutic targets (Supplementary Fig. S1B-C). As CD36 has already been shown by our and others groups to be a prognostic marker in myeloid leukemia (6,23–25), we focused on CD39. The eATPase CD39 is well known for its immunosuppressive and pro-angiogenic function in multiple cancer types (16, 17). However, its role in AML cells and its contribution to AML chemoresistance are currently unknown.

Our gene expression data indicated that CD39 expression was upregulated in residual AML cells upon AraC treatment. To confirm that enhanced transcription correlated with increased surface protein levels, we studied CD39 cell surface expression in residual viable AML cells from the bone marrow of 25 PDXs following treatment with AraC (representative flow plot in Supplementary Fig. S2A). As expected, we observed a significant cytoreduction of the total cell tumor burden in the bone marrow and spleen (Fig.1A) of these different PDX models upon AraC treatment *in vivo*. In line with our gene expression data, we observed both an increase in the percentage of CD39-positive cells and in the intensity of CD39 expression not only in the bulk residual AML population, but also in the immature CD34^+^CD38^-^ residual cell subpopulation in AraC-treated compared to vehicle-treated xenografted mice (Fig. 1B-C; Supplementary Fig. S2B-C). We then investigated the expression of CD39 in our panel of cell line-derived xenografted (CLDX) models characterized by different levels of sensitivity to AraC *in vivo (*Farge et al. 2017) and *in vitro* (representative gates strategy for *in vivo* and *in vitro* experiments in Supplementary Fig. S3A-B, respectively). AraC treatment resulted in variable induction of cell death (Supplementary Fig. S3C) and increased the transcript, as well as the cell surface expression of CD39 in our AML cells lines (HL60, MOLM14, U937 and KG1a) *in vitro* (Supplementary Fig. S3D-E). While MOLM14 and OCI-AML3 CLDX models are highly resistant to AraC chemotherapy *in vivo*, the U937 model is more sensitive and initially responds well to the treatment (total cell tumor burden fold reduction greater than 10 in AraC-*vs.* PBS-treated mice) (Fig. 1D). The majority (∼70%) of the intrinsically resistant MOLM14 and OCI-AML3 cells expressed CD39 *in vivo*. By contrast, only a small fraction (∼30%) of U937 cells expressed CD39. Interestingly, we observed a significant increase in the percentage of CD39-positive cells as well as the intensity of CD39 cell surface expression (Fig. 1 E-F), associated with an increase in CD39 eATPase activity (Fig. 1G) in residual U937 cells surviving post-chemotherapy, while no change in CD39 expression and activity was detected in MOLM-14 and OCI-AML3 cells (Fig. 1E-G). Next, we studied the kinetics of upregulation of CD39 in RLCs *in vivo,* during AraC treatment (day+3), immediately after the last dose of AraC treatment (day+5), and at day +8 in AML-xenografted NSG mice. Starting from day+5, we observed the appearance of RLCs with an increased CD39 expression (Fig. 1H). Of note, CD39-positive cells were not decreased at day 3 (Fig. 1H) and selection for CD39-positive cells occurred without genetic and mutational changes over time, as major founder mutations were present at diagnosis in patients and in the PDX throughout the same time course (Supplementary Fig. S3F). Altogether, these data strongly suggest that the CD39-positive phenotype may pre-exist before xenotransplantation and chemotherapy, and be selected and enhanced by AraC treatment *in vivo*. To test this hypothesis, we assessed whether sorted CD39^high^ and CD39^low^ cell subpopulations had a differential sensitivity to AraC treatment. Indeed, sorted CD39^high^ subsets from both MOLM14 and U937 AML cell lines pre-treated *in vitro* with AraC showed a significantly lower sensitivity to AraC with respect to their CD39^low^ counterparts (Supplementary Fig. S3G). Next, we compared *ex vivo* sensitivity to AraC of FACS-purified CD39^high^ and CD39^low^ fractions obtained from AML cells (from CLDXs and PDXs) pre-treated *in vivo* with AraC or vehicle. Strickingly, therapy-naïve AML cells expressing high levels of CD39 also exhibited a significantly higher *ex vivo* EC_50_ for AraC compared to the CD39^low^ subpopulation (Fig.1I). On the other hand, residual AML cells derived from AraC-treated mice exhibited a lower basal sensitivity to the cytotoxic drug independent of the level of CD39 expression (Fig.1I).

**Figure 1.**
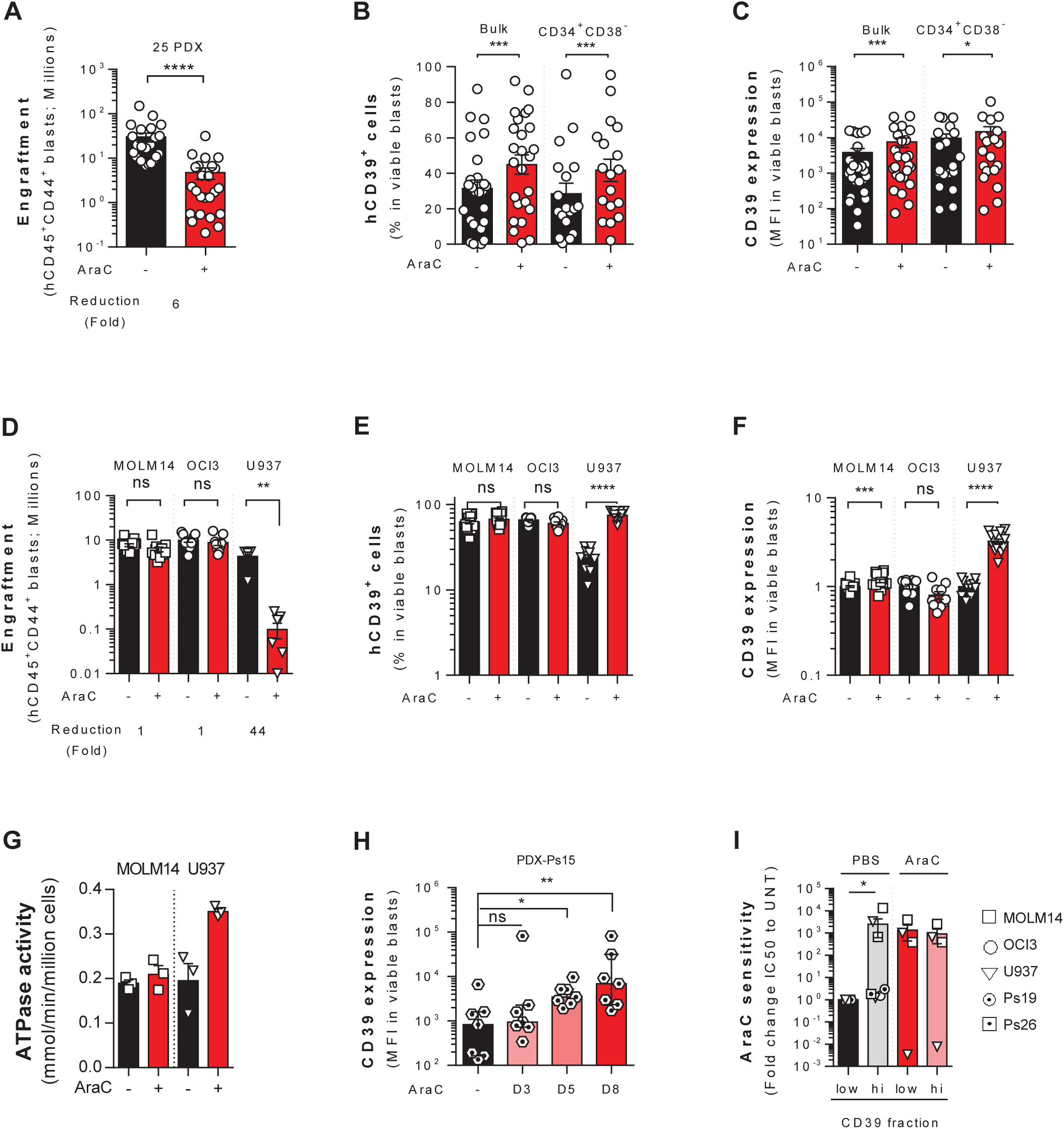
Identification of the ectonucleotidase CD39/ENTPD1 as a new actor of early resistance to cytarabine in AML. **(A)** The total number of human AML cells expressing CD45, CD33 and CD44 in 25 patient-derived xenografts (PDX) was analyzed and quantified using flow cytometry in AraC-treated compared to PBS-treated xenografted mice in bone marrow and spleen. (B-C) The percent **(B)** and MFI **(C)** of CD39^+^ cells in the bulk population and CD34^+^CD38^-^ immature cell population of human viable residual CD45^+^CD33^+^ AML cells was assessed in the bone marrow of AraC-treated compared to PBS-treated xenografted mice by flow cytometry. **(D-F)** Flow cytometry analysis of xenografts (CLDX) derived from 2 resistant AML cell lines (MOLM14, OCI-AML3) and 1 sensitive AML cell line (U937) to assess respectively **(D)** the total tumor cell burden in bone marrow and spleen of human viable AML cells in AraC- and PBS-treated CLDX, **(E)** the percentage and **(F)** MFI of CD39^+^ cells in the bone marrow of the xenografted mice. **(G)** The eATPase activity of CD39 in MOLM14 and U937 CLDX models after AraC treatment was assessed and the concentration of non-hydrolyzed extracellular ATP was determined using the ATPlite assay (PerkinElmer). **(H)** Flow cytometric analysis of human CD45^+^CD33^+^ residual AML cells derived from the bone marrow of AraC-treated AML-xenografted mice at day 3-5-8 after the start of the treatment compared to PBS-treated xenografted mice was performed to assess the expression level of CD39. **(I)** AML cell lines or primary AML samples were injected into mice, treated *in vivo* with PBS or AraC (30 mg/kg/d for CLDXs and 60 mg/kg/d for PDXs) for 5 days and sacrificed at day 8. Cells were FACS-sorted based on CD39 expression level and *ex vivo* AraC sensitivity was then evaluated. The EC50 for AraC of the sorted CD39 fractions (Low: low CD39-expressing fraction, High: high CD39-expressing fraction) is analyzed after 24h of treatment using AnnexinV/7AAD flow cytometry staining. P values were determined by the Mann-Whitney test. P-value: *≤0.05, **≤0.01, ***≤0.001, ****≤0.0001, ns= not significant.

Overall, our data indicate that a CD39^high^ phenotype characterizes *in vitro* and *in vivo* a subset of AML cells intrinsically resistant to AraC treatment. Importantly, this phenotype pre-exists and is amplified upon AraC chemotherapy *in vivo*.

### Identification of CD39/ENTPD1 as a new prognostic marker associated with poor response to chemotherapy in AML patients

In order to evaluate the clinical relevance of our findings, we analysed the expression of CD39 in AML patients. Analysis of a cohort of 162 AML patients at diagnosis indicated heterogeneous expression of CD39 (Fig. 2A). The expression of CD39 was not associated with the presence of specific recurrent mutations in AML (Supplementary Fig. S4A). However, we observed a correlation between CD39 cell surface expression and FAB classification with a lower level of expression associated with the most undifferentiated AML subtypes (Supplementary Fig. S4B). This observation was also supported by the analysis of publicly available gene expression datasets from AML patients (Supplementary Fig. S4C).

**Figure 2.**
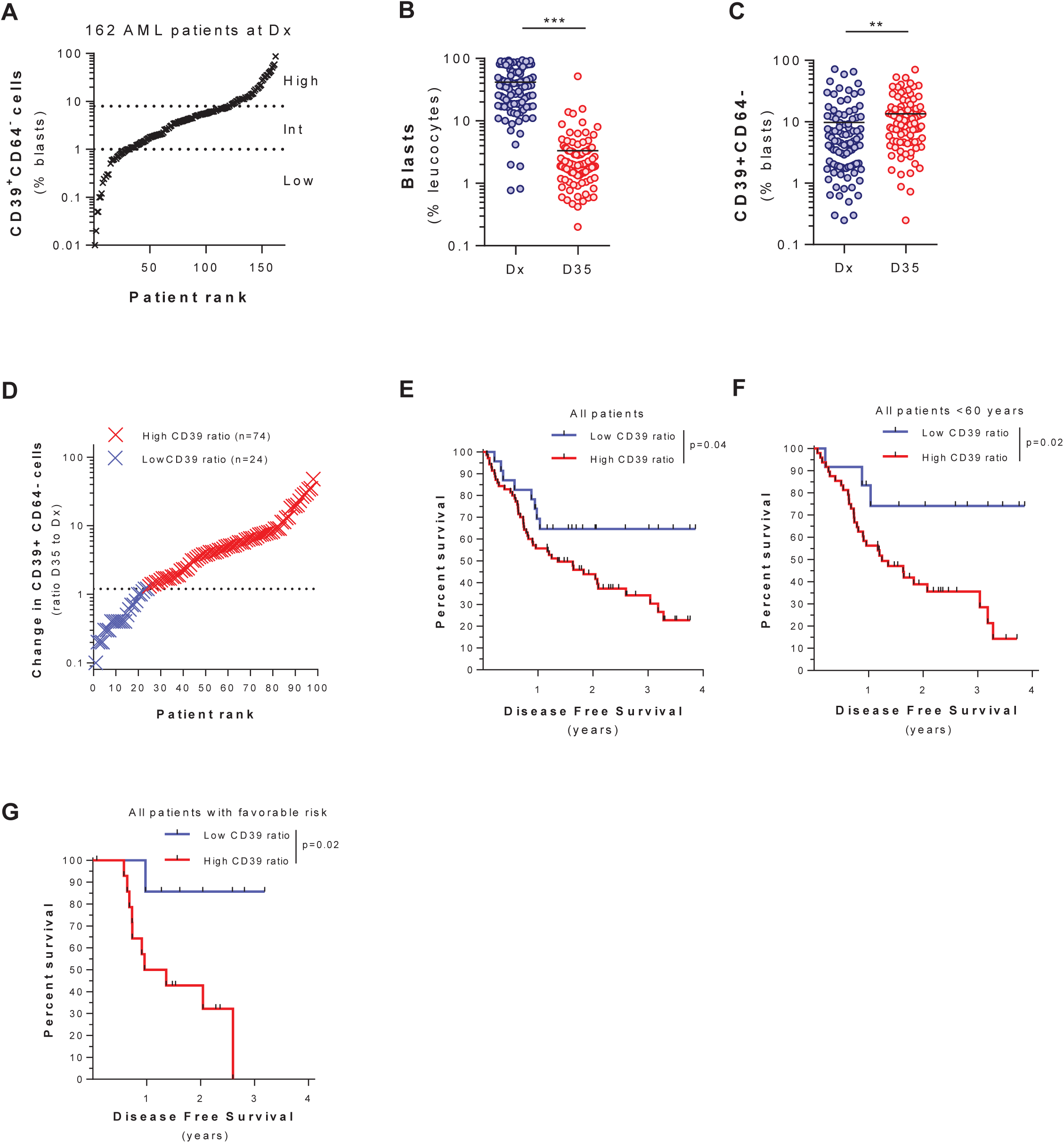
CD39 is a new prognostic marker associated with poor response to chemotherapy in AML patients. **(A)** Flow cytometric analysis of the percentage of CD39^+^ CD64^-^ blasts in the peripheral blood of 162 AML patients at diagnosis (Dx). **(B-C)** Flow cytometric analysis of the percentage of total blasts and CD39^+^CD64^+^ blasts in the peripheral blood of 98 AML patients obtained at diagnosis (Dx) and at 35 days post-chemotherapy (D35) **(B)** and distribution of the patients based on the fold-change enrichment in CD39^+^CD64^-^ blasts at day 35 vs Dx **(C)**. Patients with a fold-change >1.2 were classified as “High CD39 ratio”, while patients with a fold-change ≤1.2 were classified as “Low CD39 ratio”. **(D-F)** Disease Free Survival analysis based on the fold-change enrichment in CD39^+^CD64^-^ blasts post-chemotherapy compared to diagnosis, respectively, of the entire cohort of AML patients (n= 98; **E**), of the subgroup of patients younger then 60 years of age (n= 60; **F**) and of the subgroup of patients classified as “Favorable risk” based on the European Leukemia Net (ELN) genetic-risk classification (n= 23; **G**).

We then followed 98 of these patients comparing CD39 cell surface expression at diagnosis (Dx) and at day 35 (D35) after intensive chemotherapy. In accordance with our preclinical model, we showed a significant tumor reduction or complete remission in most of the patients after treatment (Fig. 2B), and demonstrated an overall increase in the percentage of CD39-positive cells in the residual blasts from those patients at day 35 post-intensive chemotherapy (Fig. 2C). We then stratified these AML patients based on their fold enrichment (fold>1.5) in CD39-expressing cells upon chemotherapy, defining a group of “High CD39 ratio” (n=74) and “Low CD39 ratio” patients (n=24) (Fig. 2D). Strikingly, the “High CD39 ratio” patients displayed a significantly worse disease-free survival compared to the “Low CD39 ratio” group (Fig. 2E). This survival disadvantage was even more evident when focusing on the group of patients younger than 60 years of age (Fig. 2F). Finally, we investigated whether CD39-positive cells expansion upon chemotherapy could further stratify patients classified in favorable, intermediate and high cytogenetic risk groups. The increase in CD39-positive cells did not significantly improve the prognostic classification of intermediate and high cytogenetic risk patients (Supplementary Fig. S5). However, this analysis revealed that AML patients from the favorable cytogenetic risk subgroup but characterized by a marked increase in CD39-expressing cells upon chemotherapy, displayed a significantly higher rate of short-term relapse and poorer clinical outcome (Fig. 2H).

Overall, these findings highlight the clinical relevance of our results obtained in PDXs and CLDXs and define CD39 as a marker of poor response to therapy and adverse prognosis in AML patients.

### CD39 expression is associated with with higher mitochondrial activity and biogenesis

As previous studies have demonstrated that drug-resistant AML cells exhibit high OxPHOS function and gene signatures *in vivo* (6,9,10), we investigated whether an OxPHOS gene signature was enriched in the transcriptomes of AML cells with high CD39 expression. We confirmed a positive correlation between *CD39* RNA expression and our previously defined “High OxPHOS” gene signature (6) making use of two independent transcriptomic databases from AML patients that we stratified as CD39^low^ and CD39^high^ (GSE97393: NES=-1.84, FDRq<0.001 and GSE10358: NES=-1.50, FDRq=0.004; respectively; Fig. 3A and Supplementary Fig. S6A). We then compared the metabolic status and mitochondrial activity of primary AML cells from patients with high or low levels of CD39 expression (Supplementary Fig. S6B-C). Accordingly, primary cells from CD39^high^ AML patients displayed increased CD39 eATPase activity compared to CD39^low^ patients (Fig. 3B) and this was associated with a modest increase in mitochondrial membrane potential (MMP) and larger increase in basal oxygen consumption rate (OCR, Fig. 3C-D). We then sorted the low and high CD39 cell fractions from one of our PDXs (Ps8) after treatment *in vivo* with AraC or PBS (Supplementary Fig. S6D). In line with our previous results on the primary patient samples, the *ex vivo* analysis of the metabolic status and OCR of the two cell subsets showed increased basal and maximal uncoupler-stimulated respiration, as well as ATP-linked respiration in CD39^high^ fractions compared to CD39^low^ fractions from PBS treated mice (Fig. 3E and Supplementary Fig. S6E-H). In accordance with our previously published data (6), AraC treatment resulted in the selection of residual viable AML cells with substantially increased basal and maximal uncoupler-stimulated OCR as well as ATP-linked OCR (Fig. 3E and Supplementary Fig. S6E-H). Overall, this indicated that increased levels of CD39 were associated with an enhanced mitochondrial activity and OxPHOS function in AML cells, which we previously identified as a feature of AraC resistant AML cells.

**Figure 3.**
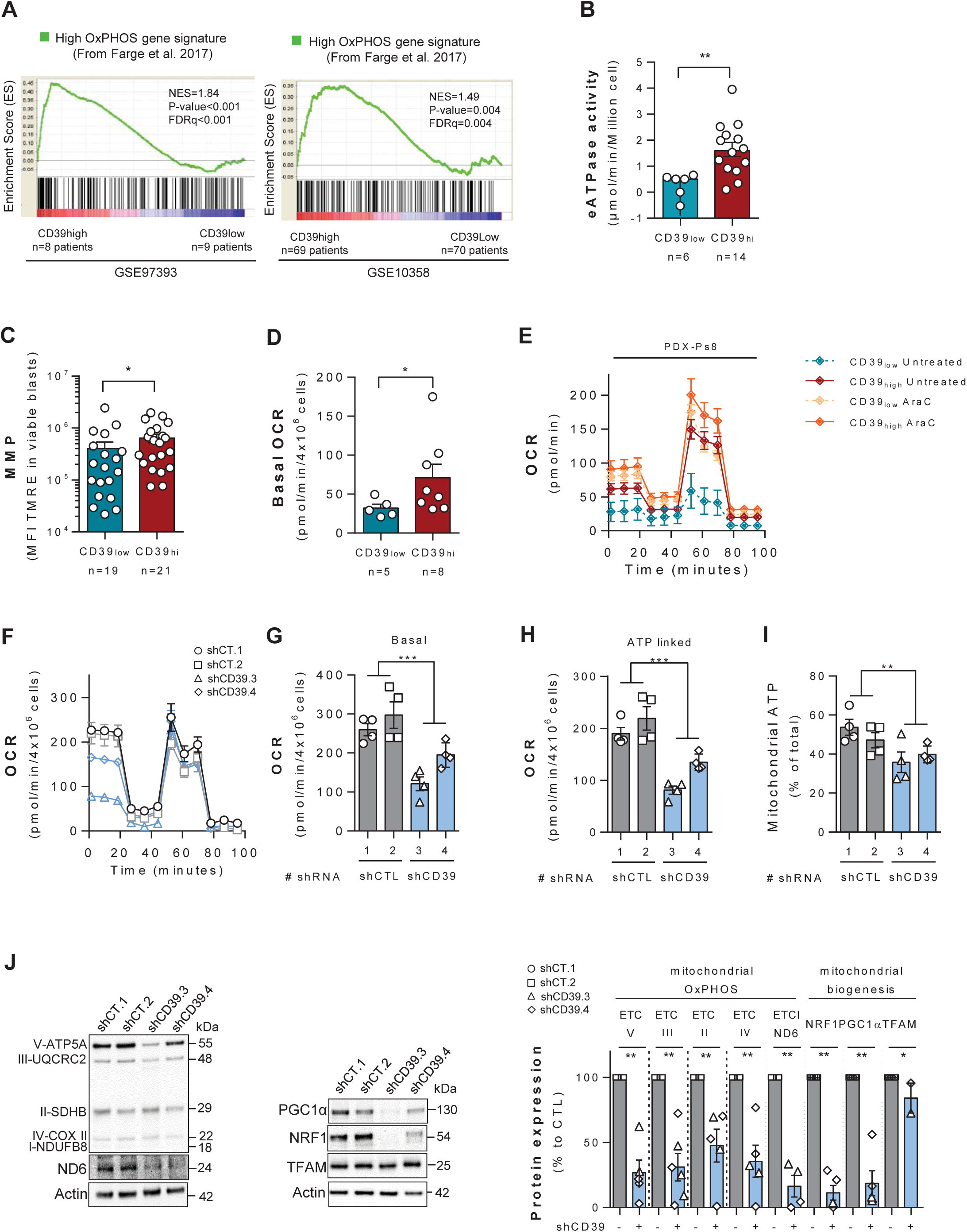
CD39 controls mitochondrial function and biogenesis. **(A)** GSEA of high OxPHOS gene signature (from Farge, 2017) was performed from transcriptomes of patients with AML classified as CD39 high *vs.* low based on the level of CD39 mRNA expression in Farge et al. 2017 (GSE97393) and TCGA cohorts (GSE10358). **(B-D)** Primary AML samples were classified based on CD39 surface expression levels and viable AML blasts were purified by FACS-sorting. Next, CD39 eATPase activity **(B)**, as well as the oxidative phosphorylation status of the CD39 high *vs.* CD39 low primary AML was analyzed *ex vivo* by measuring **(C)** the mitochondrial membrane potential using TMRE probe and **(D)** oxygen consumption rate (OCR) (basal oxygen consumption rate, and maximal oxygen consumption) assessed by Seahorse. **(E)** OCR of PDX-derived AML cells (Ps 8) obtained from leukemic mice pre-treated *in vivo* with AraC (60 mg/kg/d) or with vehicle for 5 days. CD33^+^CD44^+^ AML cells were FACS-sorted based on CD39 expression 3 days after the end of the *in vivo* treatment and their OCR was assessed *ex vivo* by Seahorse assay. **(F-I)** OCR (basal oxygen consumption rate and ATP-linked oxygen consumption) **(F-H)** and mitochondrial ATP production **(I)** assessed, respectively, using a Seahorse analyzer and Promega Cell Titer Glo kit for MOLM14 cells transduced with lentiviral vectors expressing either control (shCT.1 and shCT.2) or anti-CD39 (shCD39.3 and shCD39.4) shRNAs *in vitro*. **(J)** Protein expression of the OxPHOS mitochondrial complexes, as well as of the transcription factors PGC1α, NRF1 and TFAM was assessed by Western blot analysis in MOLM14 cells expressing anti-CD39 or control shRNAs *in vitro*. The graph on the right shows densitometric quantification of western blot bands normalized by the housekeeping gene β-Actin and relative to the average of the shCT (shCT.1 and shCT.2) samples. P-value: *≤0.05, **≤0.01, ***≤0.001.

In order to specifically study the direct effect of modulating CD39 expression on AML cell metabolism, we transduced the AML MOLM14 cell line with viral vectors expressing two different shRNAs targeting CD39. Transduction of MOLM14 with the shCD39-expressing lentiviral vectors resulted in efficient silencing of the ectonucleotidase both at the mRNA level and at the protein level (Supplementary Fig. S7A-B), leading to a significant down-regulation of the expression of this marker at the cell surface (Supplementary Fig. S7C). Silencing of CD39 resulted in a dramatic decrease in both basal and ATP-linked OCR in MOLM14 (Fig. 3F-H), which translated into a reduced generation of mitochondrial-derived ATP (Fig. 3I). This decreased mitochondrial OxPHOS activity was associated with a reduced expression of subunits of the ETC complexes and of well-known effectors of mitochondrial biogenesis (i.e. NRF1, PGC1α) (Fig. 3J). Overall, these results indicate that CD39 positively controls mitochondrial function and oxidative phosphorylation at least in part by controlling the expression of the key transcriptional activators NRF1 and PGC1α promoting mitochondrial biogenesis.

### Pharmacological inhibition of CD39 ectoducleotidase activity inhibits the metabolic reprogramming associated with AraC resistance and enhances AML cell sensitivity to AraC *in vitro*

Next, we sought to determine whether inhibition of CD39 activity by polyoxometalate 1 (POM-1), a pharmacologic inhibitor of nucleoside triphosphate diphosphohydrolase activity (26), could inhibit the metabolic reprogramming triggered by AraC and sensitize AML cells to the chemotherapic treatment *in vitro* (Fig. 4A). As expected, POM-1 inhibited the increase of the CD39 eATPase activity upon AraC treatment in all AML cell lines tested *in vitro* (MOLM14, OCI-AML3, MV4-11, each in biological triplicate; Fig. 4B), leading to an accumulation of eATP in the medium (MOLM14; Supplementary Fig. S7D). Notably, inhibition of CD39 by POM-1 in MOLM14 abrogated the expansion of the extracellular ADP and AMP pools triggered by AraC (Supplementary Fig. S7D). Furthermore, POM-1 treatment repressed the AraC-induced increase in basal OCR, mitochondrial mass, mtDNA level and the protein level of ETC subunits (Fig. 4C-F). Importantly, POM-1 treatment significantly enhanced the loss of mitochondrial membrane potential (each cell line in biological triplicate; Fig. 4G) and the induction of apoptosis (Fig. 4H) triggered by AraC treatment *in vitro* in all three AML cell lines tested.

**Figure 4.**
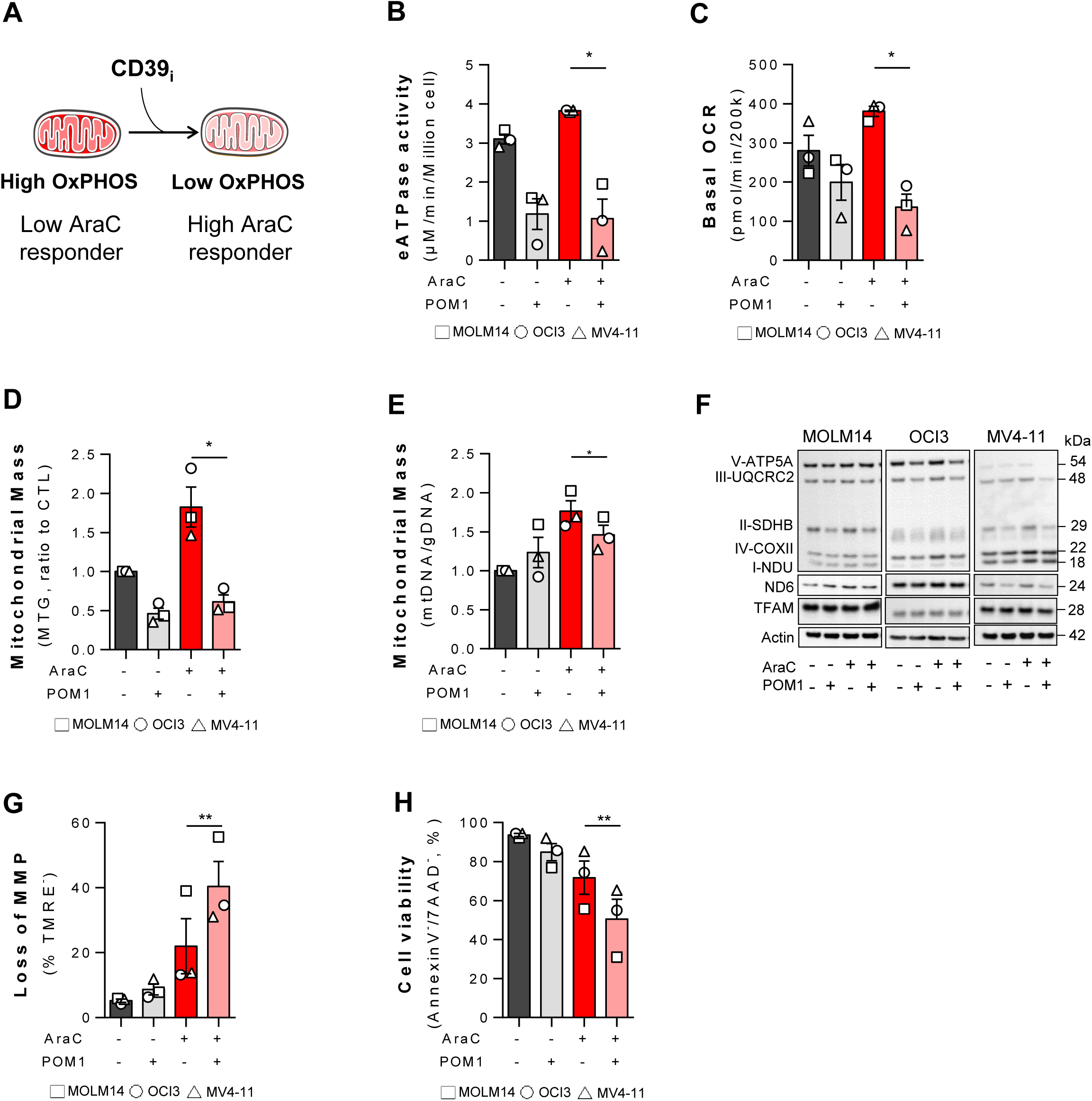
Effects of pharmacological inhibition of CD39 activity on AML metabolism and response to AraC. **(A)** Schematic depicting the metabolic reprogramming triggered by CD39 inhibition in AraC-resistant AML cells. **(B)** CD39 eATPase activity was assessed in MOLM14, OCI-AML3 and MV4-11 after 48 hours of PBS, POM-1, AraC, or POM-1+AraC treatment *in vitro*. **(C-E)** Basal OCR **(C)**, mitochondrial mass **(D)** and mitochondrial DNA content **(E)** were determined in MOLM14, OCI-AML3 and MV4-11 cultured *in vitro* for 24 hours with PBS, POM-1, AraC, or POM-1+AraC. OCR was assessed using a Seahorse analyzer **(C)**. Mitochondrial mass was assessed by flow cytometry using the fluorescent MitoTracker Green (MTG) and the values were normalized to PBS-treated samples **(D)**. Mitochondrial DNA (mtDNA) content was determined by real time PCR and the quantification was based on mtDNA to nuclear DNA (nDNA) gene encoding ratio **(E)**. **(F)** Protein expression of the mitochondrial OxPHOS complexes in MOLM14, OCI-AML3 and MV4-11 was assessed by Western blot analysis after 24-hour treatment with PBS, POM-1, AraC, or POM-1+AraC *in vitro*. **(G-H)** Loss of mitochondrial membrane potential was assessed following 48-hour treatment of MOLM14, OCI-AML3 and MV4-11 cells with PBS, AraC, POM-1, and POM-1+AraC by flow cytometry using fluorescent TMRE probe staining **(G)**. Percentage of viable cells (AnnexinV-/7AAD-) was measured after 48-hour treatment of MOLM14, OCI-AML3 and MV4-11 cells with PBS, AraC, POM-1, and POM-1+AraC by flow cytometry using AnnexinV/7-AAD staining **(H)**. P-value: *≤0.05, **≤0.01, ns=not significant. Histobars correspond to the mean of independent biological triplicates.

Altogether, our results strongly suggest that CD39 activity directly affects AML cell sensitivity to AraC through the regulation of mitochondrial function.

### Cytarabine residual cells enhance OxPHOS metabolism through the activation of the CD39-cAMP-PKA-mediated mitochondrial stress response in AML

Because our results strongly support the assertion that CD39 expression influences mitochondrial OxPHOS, we sought to explore signaling pathways downstream of CD39 that may promote OxPHOS metabolism and chemoresistance. Therefore, we performed an independent RNA expression experiment to characterize global changes induced by shCD39 in the therapy-resistant MOLM14 AML cell line. A total of 152 genes were significantly differentially expressed in MOLM14 upon silencing of CD39 (42 up-regulated, 110 down-regulated; FDR>1.25, log2(fold-change)>1.0; Fig. 5A and Supplementary Table S2). In line with our metabolic assays, gene set enrichment analysis (GSEA) indicated that CD39 loss in MOLM14 cells negatively correlated with a gene set representing High OxPHOS function (Farge et al. 2017) (NES=-1.38, FDRq=0.005; Supplementary Fig. S8A). Furthermore, the shCD39 down-regulated gene signature was significantly enriched in the transcriptomes of AML patient samples characterized by poor response to AraC *in vivo* in NSG (low *versus* high responders) (Fig. 5B) and in the transcriptomes of AraC-resistant AML cells from three AML PDXs (Fig. 5C). Gene ontology analysis of the 110 genes downregulated upon CD39 silencing (e.g. genes that also confer resistance should be positively correlated with CD39 expression) indicated an enrichment in biological processes involved in cell cycle control, DNA repair, responses to stress/stimuli, metabolism and signaling (p<0.01; Fig. 5D; Supplementary Fig. S8B). Interestingly, GSEA for known signaling pathways revealed a significant positive enrichment of genes involved in the cAMP-PKA pathway, a master regulator of mitochondrial homeostasis and oxidative stress response, and of CREB/ATF genes in transcriptomes of AML patient cells with highest CD39 expression compared to AML patient cells with lowest CD39 expression (Fig. 5E; Supplementary Fig. S6A). Moreover, transcription factor enrichment analysis identified signatures of multiple transcription factors playing key roles in mitochondrial homeostasis and stress response (such as ATF4/6, PARP1 and E2F1; Supplementary Fig. S8C). Based on these observations, we formulated the hypothesis that the activation of the cAMP-PKA-mediated stress response pathway was controlling enhanced mitochondrial activity and biogenesis driven by CD39 up-regulation upon AraC treatment in AML cells. Therefore, we investigated the modulation of the c-AMP-dependent PKA signaling pathway upon AraC treatment of MOLM14 and inhibition of CD39 (Supplementary Fig. S8E-F). Analysis of the expression and activation of key targets of the cAMP-dependent signaling pathway showed an increase in the phosphorylation of RRXS*/T*-PKA substrates upon AraC treatment (Supplementary Fig. S8D-E). These results were similar to upon activation of cAMP-PKA in MOLM14 cells treated with four diverse PKA agonists (such as FSK, IBMX, extraATP, 8-BrcAMP), while treatment with PKA antagonist (such as H89 and PKA inhibitor 14-22 Amide PKAi) inactivated cAMP-PKA pathway (Supplementary Fig. S8D). On the contrary, pharmacological and genetic inhibition of CD39 reduced intracellular cAMP and inhibited the AraC-induced phosphorylation of RRXS*/T*-PKA substrates (Supplementary Fig. S8E-F).

**Figure 5.**
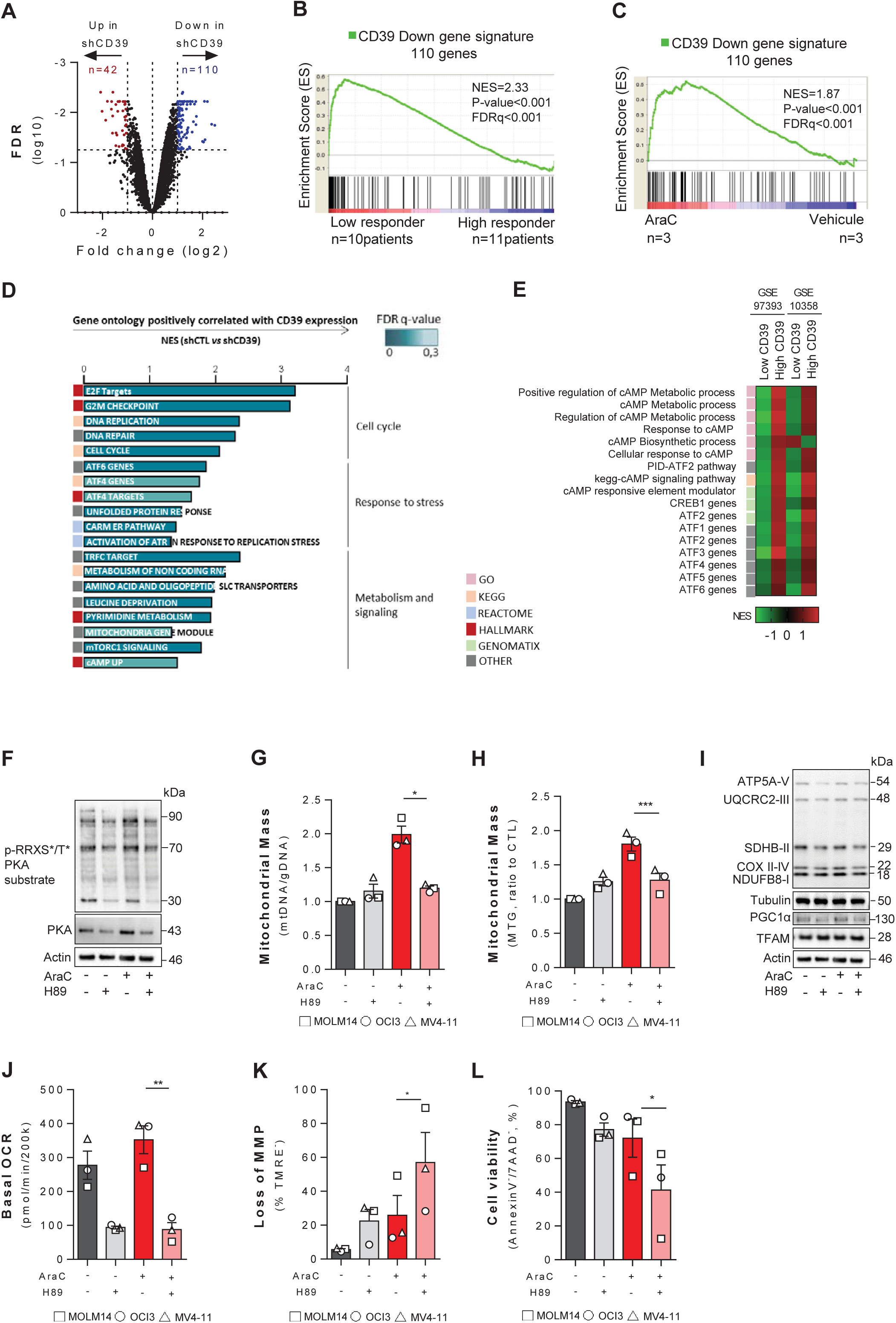
CD39 high AML chemoresistant cells maintain an enhanced OXPHOS metabolism and support mitochondrial biogenesis through the activation of a CD39-cAMP-PKA axis. **(A)** Volcano plot displaying differential expressed genes between MOLM14 silenced for CD39 expression (shCD39) and control cells (shCT). On the y-axis the FDR values (log10) are plotted, while the x-axis displays the fold-change (FC) values (log 2). The red dots represent the upregulated (FC>-1.0, FDR>1.25), while the blue dots represent the down-regulated (FC>1.0, FDR>-1.25) expressed transcripts in shCD39 *vs.* shCT MOLM14. **(B-C)** GSEA of the shCD39 down-regulated gene signature (n= 110 genes) was performed from **(B)** the transcriptomes of AML patient samples characterized by poor (red) compared to good (blue) response to AraC *in vivo* upon xenotransplantation in NSG mice (low *vs*. high responders) and **(C)** from the transcriptomes of human residual AML cells purified from AraC-treated (red) compared with vehicle (PBS)-treated (blue) AML-xenografted NSG mice (AraC *vs*. Vehicule). **(D)** Gene set enrichment analysis (GSEA) of the AML cell line MOLM14 shCT *versus* shCD39. Positively enriched gene ontology terms. Bar length, normalized enrichment score (NES). Bar color, FDR. **(E)** GSEA of signaling pathways gene signature was performed from transcriptomes of patients with AML that had the highest CD39 mRNA expression compared with those with the lowest expression in Farge (2017) and TCGA cohorts. **(F)** Protein expression of phospho-PKA substrate (RRXS*/T*) and PKA was assessed by Western blot analysis after 6-hour treatment with AraC and/or H89 in MOLM14 cells *in vitro*. The housekeeping gene β-Actin was used as loading control. **(G)** Mitochondrial DNA content was determined in MOLM14, OCI-AML3 and MV4-11 upon treatment *in vitro* for 24-hours with PBS, H89, AraC, or H89+AraC by real time PCR. Quantification was based on mtDNA to nuclear DNA (nDNA) gene encoding ratio. **(H)** Mitochondrial mass was assessed by flow cytometry using the fluorescent MitoTracker Green (MTG), in MOLM14, OCI-AML3 and MV4-11 cells after PBS, H89, AraC, or H89+AraC 24-hour treatment. The values were normalized to PBS-treated samples. **(I)** Protein expression of mitochondrial OxPHOS complexes and PGC1α was assessed by Western blot analysis after 24-hour treatment with AraC and/or H89 in MOLM14 cells *in vitro*. The housekeeping gene α-Tubulin and actin were used as loading control. **(J)** Basal OCR in MOLM14, OCI-AML3 and MV4-11 cells after PBS, H89, AraC, or H89+AraC 24-hour treatment was assessed using a Seahorse analyzer. **(K-L)** MOLM14, OCI-AML3 and MV4-11 cells was treated with PBS, AraC, H89, and H89+AraC for 48-hour to assess **(K)** loss of mitochondrial membrane potential by fluorescent TMRE probe and **(L)** percentage of viable cells using Annexin V/7-AAD staining by flow cytometry. P-value: *≤0.05, **≤0.01, ***≤0.001, ns=not significant. Histobars correspond to the mean of independent biological triplicates..

We next investigated whether inactivation of PKA pathway by H89, a well-known pharmacological agent that inhibits PKA activity, could affect AML mitochondrial functions and enhance AraC treatment cytotoxicity similarly to CD39 inhibition. As expected, H89-treated MOLM14 cells exhibited decreased levels of p-RRXS*/T*-PKA substrates, including in AraC setting (Fig. 5F). Importantly, the PKA-specific inhibitor H89 significantly phenocopied the effect of the CD39 inhibitor POM-1 on mitochondrial activity by counteracting the increase in mtDNA level (Fig. 5G), mitochondrial mass (Fig. 5H), the expression of ETC complex subunits and PGC1α (Fig. 5I), and basal OCR (Fig. 5J) induced by AraC treatment. Finally, H89-treated MOLM14 cells exhibited an increased loss of MMP (Fig. 5K) and a reduction of cell viability (Fig. 5L).

These results strongly support the hypothesis that CD39 activity greatly influences AraC cytotoxicity through modulation of mitochondrial function in a cAMP-PKA-dependent manner.

### Targeting CD39 enhances AraC chemotherapy efficacy *in vivo*

Our data suggest that inhibiting CD39 activity may be a promising therapeutic strategy to enhance chemotherapy response of AML cells *in vivo*. In order to test this hypothesis, we generated NSG mice-based CLDX and PDX models from AML cell lines and primary patient cells, respectively. We then tested the consequences of CD39 repression on the response of our pre-clinical models of AML to AraC using as alternative experimental strategies genetic invalidation and pharmacological inhibition of our target. Genetic invalidation of CD39 was achieved in the MOLM14 CLDX model *in vivo* using two different doxycycline-inducible shRNAs specifically targeting the *ENTPD1* gene. Remarkably, CD39 depletion in combination with AraC treatment resulted in a significant reduction of total cell tumor burden in the bone marrow of the mice three days after the end of the chemotherapy cycle (day 18) compared to the both vehicle treated counterparts and the shCTL leukemias (Fig. 6A-B). Moreover, while no change in AML viability and loss of MMP was detectable in control MOLM14 upon AraC treatment, concomitant repression of CD39 triggered a significant decrease in viability and loss of MMP in AML cells (Fig. 6C-D). Altogether, this led to an enhanced AraC sensitivity *in vivo* as further demonstrated by a significant increase in the overall survival of AraC-treated shCD39-xenografted mice compared to both the vehicle-treated shCD39-xenografted mice cohort and the AraC-treated shCTL-xenografted mice (Fig. 6E).

**Figure 6.**
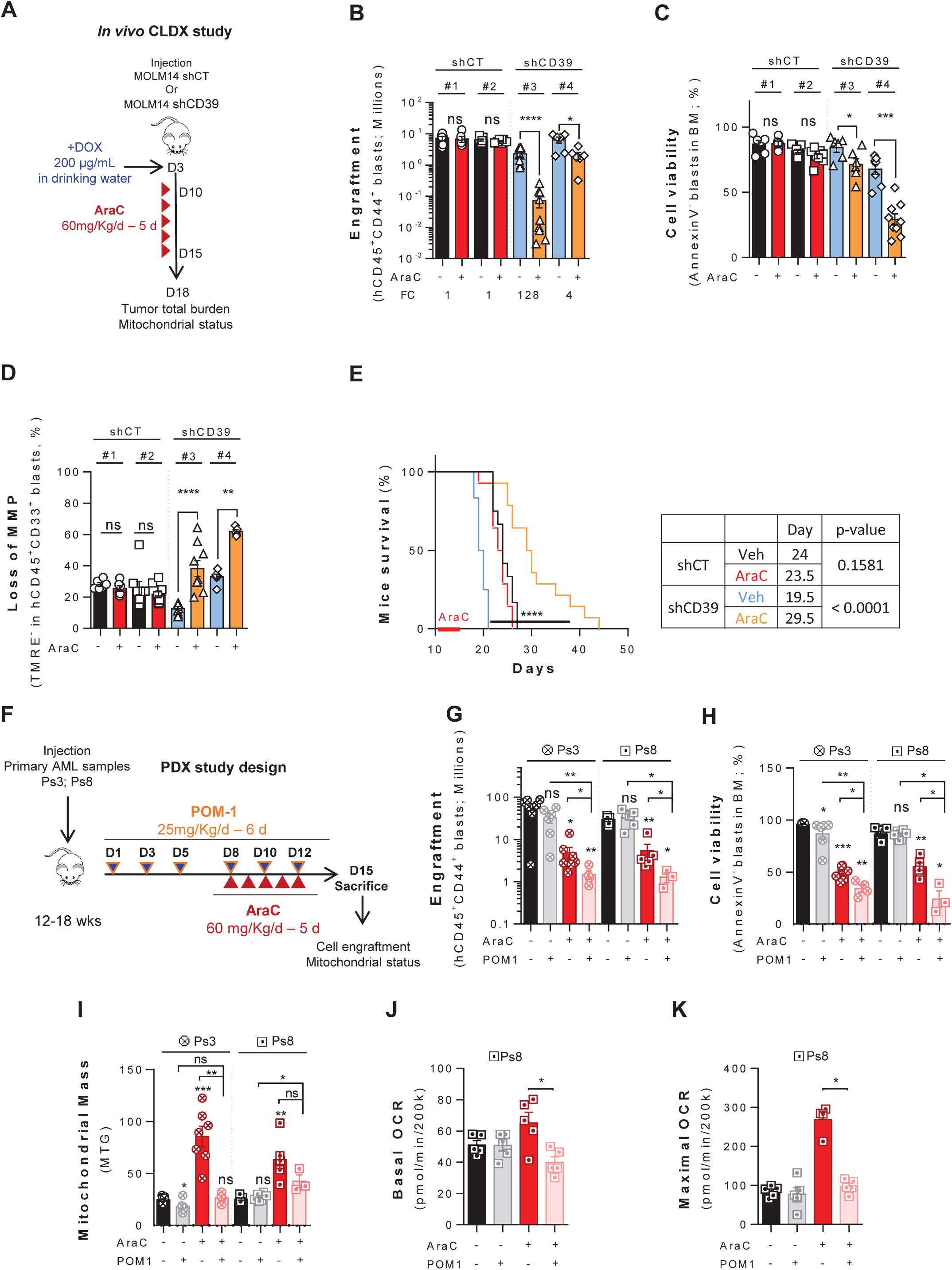
*In vivo* targeting CD39 sensitizes to cytarabine in AML-engrafted mice. **(A)** Schematic diagram of the chemotherapy regiment and doxycycline-administration schedule used to treat CLDX NSG mice xenografted with MOLM14 transduced with shCT or shCD39 shRNA-expressing lentiviral vectors. The CLDX (shCTL MOLM14, shCD39 MOLM14) models were treated with vehicle (PBS) or 60 mg/kg/day AraC given daily *via* intraperitoneal injection for 5 days. Mice were sacrificed post-treatment at day 15 and AML cells were harvested for analysis. **(B)** Cumulative total cell tumor burden of human viable CD45^+^CD44^+^CD33^+^ AML cells transduced with shCD39 or shCT was assessed in bone marrow and spleen for the indicated groups of mice by flow cytometry**. (C-D)** Percent of human viable AML cells **(C)** and loss of mitochondrial membrane potential **(D)** in AML cells in the bone marrow of the leukemic mice were assessed by flow cytometry using AnnexinV/7-AAD and fluorescent TMRE probe staining, respectively. **(E)** The overall-survival of the mice transplanted with shCTL or shCD39 MOLM14 cells and treated with AraC or vehicle as described in **(A)** is shown. **(F)** Schematic diagram of the chemotherapy regimen and schedule used to treat NSG-based PDX (Ps3, Ps8) models with vehicle, AraC or POM-1 CD39 inhibitor or with a combination of the latter two. Mice were treated with the CD39 inhibitor (25 mg/kg/day) every other day for two weeks. In parallel, mice were treated with vehicle (PBS) or 60 mg/kg/day AraC given daily *via* intraperitoneal injection for 5 days. Mice were sacrificed post-treatment at day 15 and the tumor total burden and the oxidative and mitochondrial status in viable AML cells was assessed. **(G-H)** Total cell tumor burden of human viable CD45^+^CD44^+^CD33^+^ AML in the bone marrow and spleen **(G)**, as well as the percentage of viable human CD45^+^CD33^+^ AML cells in the bone marrow **(H)** was assessed at day 15, 3 days after the end of the therapy. **(I-K)** The Mitochondrial mass **(I)** was assessed by flow cytometry using the fluorescent probe MitoTracker Green (MTG) and, the oxygen consumption rate (basal and maximal OCR; **J-K**) was assessed by seahorse in FACS-sorted human AML cells harvested From the mice at day 15, 3 days after the end of the therapy. P-value: *≤0.05, **≤0.01, ***≤0.001, ****≤0.0001, ns= not significant.

Pharmacological inhibition of CD39 activity was also achieved by administering the eATPase inhibitor POM-1 for 6 days at the dose of 25 mg/kg/day alone and in combination with AraC at 60 mg/kg/day for 5 consecutive days in two different and independent PDX models (Ps3, Ps8). Response to single and combinatory treatments and various characteristics of RLCs were specifically monitored at day 15 (3 days after the last administration of the combinatorial treatment; Fig. 6F). Similar to our previous results, we observed an enhancement of AraC cytotoxic effect in combination with POM-1 administration in two PDXs with a significant reduction of the total cell tumor burden and of AML cell viability in the bone marrow *in vivo* in the combination treatment group compared to both the single treatment groups (Fig. 6G-H). We confirmed that POM-1 efficiently blocked *in vivo* the CD39 eATPase activity induced upon AraC treatment in these different 2 PDX models (Ps3, Ps8; Supplementary Fig. S9A) and induced a significant reduction of the mitochondrial mass (Fig. 6I), and basal and maximal mitochondrial respiration upon the combination treatment compared to AraC alone (Fig. 6J-K and Supplementary Fig. S9B).

In summary, our results indicate that AraC treatment induces the selection and amplification of CD39 expressing pre-existent and intrinsically resistant AML cells. CD39 activity promotes AraC resistance by activating cAMP-mediated mitochondrial stress response leading to the induction of the expression of key master regulators of mitochondrial biogenesis and homeostasis that modulate mitochondrial OxPHOS function in RLCs (Fig. 7). Increase in CD39 expression is associated with poor prognosis in the clinics. Furthermore, our results show that inhibition of CD39 expression or activity substantially improves the response to cytarabine treatment in preclinical models of AML *in vivo*. Therefore, our findings strongly support the rationale for targeting CD39 as a valuable therapeutic strategy to enhance response to AraC in therapy resistant AML. This should be assessed through a clinical study for AML treatment combining anti-CD39 small molecules with cytarabine chemotherapy.

**Figure 7.**
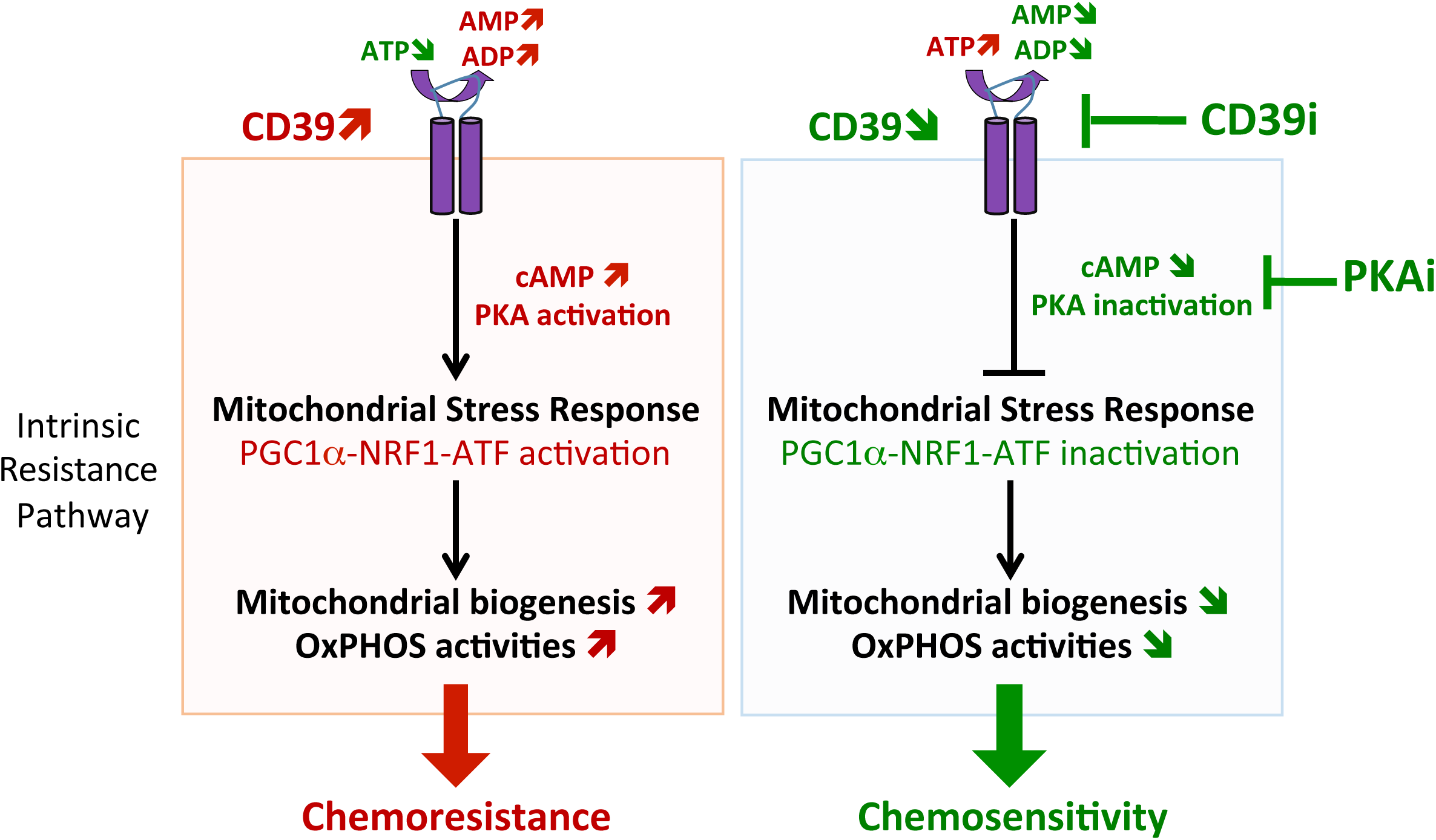
CD39-cAMP-PKA-mediated mitochondrial and metabolic reprogramming is involved in the resistance to AraC in AML. Schematic diagram of AraC mechanism of resistance involving the CD39-dependent crosstalk between energetic niche and AML mitochondrial functions through CD39-cAMP-PKA signaling axis. Intrinsic PKA pathway through PGC1a supports mitochondrial biogenesis to maintain high OxPHOS metabolism, cell survival and chemoresistance upon AraC treatment.

## DISCUSSION

Poor overall survival is mainly due to frequent relapse caused by RLCs in AML patients (2, 27). While recent studies have highlighted new mechanisms of drug resistance in AML especially *in vivo* (6,28,29), their clinical applications are still unresolved or under assessment. New therapies that specifically target and effectively eradicate RLCs represent an urgent medical need. In this work, we have identified the cell surface eATPase ENTPD1/CD39 and its downstream signaling pathway as a new critical and druggable target involved in the resistance to cytarabine in AML. We showed that CD39 was overexpressed in residual AML cells post-chemotherapy from both 25 PDX models and 98 patients in the clinical setting. CD39 is also highly expressed in several human solid tumors, in which it was shown to actively contribute to cancer cell proliferation, dissemination and metastatic process (30, 31). In the context of AML, our data supports a model in which AraC treatment induces the selection and amplification of CD39 expressing pre-existent and intrinsically resistant leukemic cells. We have furthermore observed that drug-induced increase in CD39 expression is associated with a poor response to AraC *in vivo*, persistence of residual disease, and with poor overall survival in AML patients, especially in the younger subgroup of patients.

While many pro-survival and anti-apoptotic signals are activated in AML by the stroma (32, 33), nucleotides and nucleosides have emerged as important modulators of tumor biology. In particular, ATP and adenosine are major signaling molecules present in the tumor microenvironment. A growing body of evidence shows that when these molecules are released by cancer cells or surrounding tissues, they act as prometastatic factors, favoring tumor cell migration and tissue colonization. Interestingly, eATP elicits different responses in tumor cells, including cell proliferation (34, 35), cell death (36, 37) and metastasis (38, 39). Furthermore, several studies described a direct cytotoxicity of eATP on different tumor cell types such as melanoma, glioma and colon cancer cells (40–42). In AML, eATP was reported to reduce human leukemia growth *in vivo* and enhance the antileukemic activity of AraC (22). The increase of CD39 expression occurred at early time points of the chemotherapeutic response and residual disease processes in PDXs and patients likely due to enrichment in eATP released from dying and apoptotic cells upon AraC treatment. Of note, high expression of CD39 was associated with a higher activity resulting in eATP hydrolysis to support AML regrowth and relapse. Bone marrow microenvironment is a key regulator of leukemia growth and has many chemoprotecting effects for AML cells (23,43,44). We and others have shown that mitochondrial OxPHOS is a crucial contributing factor of AML chemoresistance and its inhibition sensitizes cells to AraC treatment (6,7,45). This is mainly due to an increase in respiratory substrate availability and in mitochondrial machinery transfer from the BM-MSCs (46, 47). Here we unveil an additional mechanism supporting mitochondrial high OxPHOS activity in reponse to AraC treatment through the enhancement of mitochondrial biogenesis triggered by CD39 eATPase activity and the downstream activation of cAMP-PKA signaling. Indeed, upon AraC-induced CD39 up-regulation, we observed an increased replication of mtDNA and expression of PGC-1α and NRF1, well known central transcriptional regulators of mitochondrial biogenesis (Fig. 7) (48, 49). Conversely, inhibition of CD39 or of the cAMP-PKA pathway led to the inhibition of mitochondrial biogenesis and OxPHOS activity increasing the cytotoxicity of AraC treatment in AML. Furthermore, our results indicate that reversible activation of the PKA pathway upon CD39 blockade and AraC treatment similarly promotes PGC1α-induced mitochondrial biogenesis and stimulates OxPHOS metabolism as reported in several other recent works (50, 51).

Previous studies have reported the pleiotropic roles of cAMP signaling and its major downstream effector PKA in different cancers including AML. Perez and colleagues showed that cAMP efflux from the cytoplasm protects AML cells from apoptosis (52). Similarly, others reported cAMP mediated protection of acute promyelocytic leukemia against anthracycline (53) or against arsenic trioxide-induced apoptosis (54). PKA, whose activation initiates an array of transcriptional cascades involved in the immune response, cell metabolism and mitochondrial biogenesis, is one of the main and canonical downstream effectors of cAMP signaling. Intriguingly, cAMP-PKA signaling can be localized not only on the plasma membrane or nucleus but also on the outer mitochondrial membrane or matrix (55). Mitochondrial cAMP signaling was shown to regulate cytosol-mitochondrial crosstalk, mitochondrial biogenesis and morphology, mitochondrial dynamics, mitochondrial membrane potential, TCA cycle activities and ETC complexes in basal and stress conditions such as starvation or hypoxia (56). Of note, cAMP signaling is activated as integral part of the mitochondrial stress response that allows the rewiring of cellular metabolism in the presence of cellular damage and oxidative stress conditions. In this context, PGC1α and NRF1/2 activation leads to an increase in expression and assembly of respiratory chain supercomplexes and a boost in oxidative phosphorylation activity, allowing dynamic adaptation of mitochondrial functions to survive adverse conditions (57). These studies establish a mechanistic link between cAMP, PKA and PGC1α in the regulation of mitochondrial biogenesis/function through the activation of mitochondrial stress response.

Collectively, our data suggest that CD39 activity, through the control of extracellular levels of ADP and AMP and the downstream activation of the cAMP-PKA pathway, may trigger a process similar to the mitochondrial stress response in resistant leukemic cells to rewire their energetic metabolism and enhance PGC1α-mediated mitochondrial biogenesis and OxPHOS activity upon chemotherapy treatment (Fig. 7). In this context, we propose that eATP and CD39 are key actors in a novel signaling mechanism implicated in AML chemoresistance to AraC and that targeting CD39 would be a promising therapeutic strategy to sensitize AML cells to AraC. In light of the recently recognized “immune checkpoint mediator” function of CD39 that interferes with anti-tumor immune responses, our data further suggest the existence of a critical crosstalk between AML cells and their immune and stromal microenvironment mediated by extracellular nucleotides and/or CD39 in the response to therapy of AML cells. In this context, blocking CD39 activity could have a double edge therapeutic benefit by both dampening the metabolic reprogramming supporting AraC cell-autonomous resistance and disrupting the immune escape mechanisms.

In conclusion, our study uncovers a non-canonical role of CD39 on AML resistance (Fig.7), and provides a strong scientific rationale for testing CD39 blockade strategies in combination with AraC treatment in clinical trials for patients with AML. Because CD39-blocking monoclonal antibodies are already in clinical trials as a single agent and in combination with an approved anti-PD-1 immunotherapy or standard chemotherapies for patients with lymphoma or solid tumor malignancies, we expect that these findings have the potential for rapid translation of our proposed combination therapy with CD39 as a putative predictive biomarker into the clinic.

## METHODS

### Primary cells from AML patients

Primary AML patient specimens are from Toulouse University Hospital (TUH, Toulouse, France). Frozen samples were obtained from patients diagnosed with AML at TUH after signed informed consent in accordance with the Declaration of Helsinki, and stored at the HIMIP collection (BB-0033-00060). According to the French law, HIMIP biobank collections have been declared to the Ministry of Higher Education and Research (DC 2008-307 collection 1) and obtained a transfer agreement (AC 2008-129) after approval by the “Comité de Protection des Personnes Sud-Ouest et Outremer II” (ethical committee). Clinical and biological annotations of the samples have been declared to the CNIL (“Comité National Informatique et Libertés”; i.e. “Data processing and Liberties National Committee”). Peripheral blood and bone marrow samples were frozen in fetal calf serum with 10% DMSO and stored in liquid nitrogen. The percentage of blasts was determined by flow cytometry and morphological characteristics before purification.

### AML mouse xenograft model

Animals were used in accordance to a protocol reviewed and approved by the Institutional Animal Care and Use Committee of Région Midi-Pyrénées (France). NOD/LtSz-scid/IL-2Rγchain^null^ (NSG) mice were produced at the Genotoul Anexplo platform at Toulouse (France) using breeders obtained from Charles River Laboratory. Mice were housed and human primary AML cells were transplanted as reported previously (58–60). Briefly, mice were housed in sterile conditions using HEPA filtered micro-isolators and fed with irradiated food and sterile water. Transplanted mice were treated with antibiotic (baytril) for the duration of the experiment. Mice (6-9 weeks old) were sublethally treated with busulfan (30 mg/kg/d) 24hr before injection of leukemic cells. Leukemia samples were thawed at room temperature, washed twice in PBS, and suspended in Hanks balanced salt solution at a final concentration of 1–10 million cells per 200 µL of Hanks balanced salt solution per mouse for tail vein injection. Daily monitoring of mice for symptoms of disease (ruffled coat, hunched back, weakness and reduced mobility) determined the time of killing for injected animals with signs of distress. If no signs of distress were seen, mice were initially analyzed for engraftment 8 weeks after injection except where otherwise noted.

### Cytarabine treatment in vivo

8 to 18 weeks after AML cell transplantation and successful engraftment in the mice (tested by flow cytometry on peripheral blood or bone marrow aspirates), NSG mice were treated by daily intraperitoneal (IP) injection for 5 days of 30 (for CLDX models) and 60 (for PDX models) mg/kg AraC, kindly provided by the Pharmacy of the TUH (Toulouse, France). For control, NSG mice were treated daily with IP injection of vehicle, PBS 1X. Mice were monitored for toxicity and provided nutritional supplements as needed.

POM-1 or ARL67156 was administrated to xenografted mice by IP injection every other day for two weeks. The time of dissection was fifteen days after the last dose of POM-1 (or ARL67156) or 8 days for AraC, two days after the last dose of each treatment.

### Assessment of leukemic engraftment

NSG mice were humanely killed in accordance with European ethic protocols. Bone marrow (mixed from tibias and femurs) and spleen were dissected in a sterile environment and flushed in Hanks balanced salt solution with 1% FBS, washed in PBS and dissociated into single cell suspensions for analysis by flow cytometry of human leukemic cell engraftment and bone marrow cell tumor burden. MNCs from peripheral blood, bone marrow and spleen were labeled with FITC-conjugated anti-hCD3, PE-conjugated anti-hCD33, PerCP-Cy5.5-conjugated anti-mCD45.1, APCH7-conjugated anti-hCD45 and PeCy7-conjugated anti-hCD44 (all antibodies from Becton Dickinson, BD, except FITC-conjugated anti-hCD3 from Ozyme Biolegend) to determine the fraction of human blasts (hCD45^+^mCD45.1^-^hCD33^+^hCD44^+^ cells) using flow cytometry. Analyses were performed on a Life Science Research II (LSR II) flow cytometer with DIVA software (BD) or Cytoflex flow cytometer with CytoExpert software (Beckman Coulter). The number of AML cells/ul peripheral blood and the cumulative number of AML cells in bone marrow and spleen (total tumor cell burden) were determined by using CountBright beads (Invitrogen) using previously described protocols (Sarry et al 2011, Farge et al 2017).

### Cell lines and culture conditions

Human AML cell lines were maintained in Roswell Park Memorial Institute (RPMI) 1640 supplemented with 10% fetal bovine serum (Invitrogen, Carlsbad, CA, USA) in the presence of 100 units per ml of penicillin and 100 μg/ml of streptomycin, and were incubated at 37°C with 5% CO_2_. The cultured cells were split every 2–3 days and maintained in an exponential growth phase. U937 was obtained from the DSMZ (Braunschweig, Germany) in February 2012 and from the ATCC (Manassas, VA, USA) in January 2014. MV4-11 and HL-60 were obtained from the DSMZ in February 2012 and February 2016. KG1 was obtained from the DSMZ in February 2012 and from the ATCC in March 2013. KG1a was obtained from the DSMZ in February 2016. MOLM14 was obtained from Pr. Martin Carroll (University of Pennsylvania, Philadelphia, USA) in 2011 and from the DSMZ in June 2015. DSMZ and ATCC cell banks provides authenticated cell lines by cytochrome C oxidase I gene (COI) analysis and short tandem repeat (STR) profiling. Furthermore, the mutation status was also verified by targeted re-sequencing of a panel of 40 genes frequently mutated in AML as described in Supplementary methods. Clinical and mutational features of our AML cell lines are described in Supplementary Table S1. These cell lines have been routinely tested for *Mycoplasma* contamination in the laboratory.

### Statistical analyses

We assessed the statistical analysis of the difference between 2 sets of data using non-parametric Mann-Whitney test one-way or two-way (GraphPad Prism, GraphPad). The Mantel-Cox log-rank test was used for statistical assessment of survival. *P* values of less than 0.05 were considered to be significant (* *P*<0.05, ** *P*<0.01 and *** *P*<0.001).

For *in vitro* and *in vivo* analyses of cytarabine residual disease and CD39 studies, see Supplementary Methods. RNA-seq data are available at the Gene Expression Omnibus under the accession number GSE136551.

## Acknowledgements

We thank all members of mice core facilities (UMS006, ANEXPLO, Inserm, Toulouse) in particular Marie Lulka, Katia Pilipenko, Christine Campi, Pauline Challies, Pauline Debas, Yara Bareira for their support and technical assistance, Cécile Déjou (Institut de Recherche en Cacérologie de Montpellier) for her help with CD39 activity assays and Prof. Véronique De Mas and Eric Delabesse for the management of the Biobank BRC-HIMIP (Biological Resources Centres-INSERM Midi-Pyrénées “Cytothèque des hémopathies malignes”) that is supported by CAPTOR (Cancer Pharmacology of Toulouse-Oncopole and Région). We thank Anne-Marie Benot, Muriel Serthelon and Stéphanie Nevouet for their daily help about the administrative and financial management of our Team. We also thank the patients and the Association GAEL for their generous support. The authors also thank Dr Mary Selak for critical reading of the manuscript.

